# The effect of breaking up sedentary time with calisthenics on neuromuscular function: a preliminary study

**DOI:** 10.1101/2022.06.22.497167

**Authors:** Emily Mear, Valerie Gladwell, Jamie Pethick

**Author notes:** Corresponding author: Dr Jamie Pethick, School of Sport, Rehabilitation and Exercise Sciences, University of Essex, Wivenhoe Park, Colchester, CO4 3WA, United Kingdom.

## Abstract

Prolonged sedentary behaviour has a detrimental effect on neuromuscular function and is associated with decreased muscle strength and force control, and a decreased ability to maintain balance. Breaking up sedentary time with regular bouts of physical activity has numerous health benefits, though the effects on neuromuscular function are unknown. This study aimed to investigate the effect of breaking up sedentary time with calisthenic exercise on neuromuscular function. To that end, 17 healthy adults, who spent ≥6 hours a day sitting, were randomly assigned to a four-week calisthenics intervention (n = 8) or a control group (n = 9). The calisthenics intervention involved performing up to eight sets of exercises during the working day (09:00-17:00); with one set consisting of eight repetitions of five different exercises (including squats and lunges). Before and immediately after the intervention, measures of knee extensor maximal voluntary contraction (MVC; right leg only) and submaximal force control (measures of the magnitude and complexity of force fluctuations; right leg only), and dynamic balance (Y balance test; both legs) were taken. The calisthenics intervention resulted in a significant increase in knee extensor MVC (*P* = 0.036), significant decreases in the standard deviation (*P* 0.031) and coefficient of variation (*P* = 0.016) of knee extensor force fluctuations during contractions at 40% MVC, and a significant increase in Y balance test posterolateral reach with left leg stance (*P* = 0.046). These results suggest that breaking up sedentary time with calisthenics may be effective at increasing muscle strength, force steadiness and dynamic balance.

**New findings:** *What is the central question of this study?:* This study sought to determine whether breaking up sedentary time with a (4-week) calisthenics exercise intervention could improve neuromuscular function.

*What is the main finding and its importance?:* A 4-week calisthenic exercise intervention increased knee extensor maximal strength, knee extensor force steadiness during submaximal contractions, and aspects of dynamic balance. These results indicate the regularly breaking up sedentary time with calisthenics can mitigate against the negative effects of prolonged sedentary time.

## Introduction

It has been estimated that adults spend 51-68% of their waking time in sedentary behaviours (Dunstan *et al.*, 2012), defined as activities done while sitting or reclining with energy expenditure ≤ 1.5 METs (Tremblay *et al.*, 2017). Indeed, technological and social factors have made sitting the most ubiquitous behaviour during working, domestic and recreational time (Healy *et al.*, 2009; Parry *et al.*, 2012). Moreover, evidence from around the world suggests that time spent in sedentary behaviours increased by ~20% as a result of COVID-19 lockdowns (Castañeda-Babarro *et al.*, 2020; Romero-Blanco *et al.*, 2020). Sedentary behaviour, and a consequent lack of muscle contractile activity, drives numerous maladaptive metabolic and cardiovascular responses (Hamilton *et al.*, 2007; Hamilton, 2018). Worryingly, time spent in sedentary behaviours is associated with increased risk of at least 35 pathological and clinical conditions (Booth *et al.*, 2012) and all-cause mortality (Lee *et al.*, 2012), even after accounting for the effects of participating in recommended amounts of moderate and vigorous physical activity (Koster *et al.*, 2012; Ekelund *et al.*, 2016).

Sedentary behaviour also has a profound effect on neuromuscular physiology and, consequently, function. Experimentally induced periods of muscle disuse, brought about by short-term limb immobilisation (a more extreme form of sedentary behaviour), have been demonstrated to decrease muscle cross-sectional area (Adams *et al.*, 1994), decrease the ability to voluntarily activate motor units (Clark *et al.*, 2006) and decrease motor unit firing rates (Duchateau and Hainaut, 1990; Seki *et al.*, 2001). These changes likely mediate the observed negative changes in neuromuscular output, and ability to perform activities of daily living (ADLs), seen with sedentary behaviour. Observational studies have demonstrated that greater time spent in sedentary behaviour is negatively associated with muscle strength (Volkers *et al.*, 2012; Hamer and Stamatakis, 2013), whilst limb immobilisation studies have demonstrated decreased muscle force control during submaximal isometric contractions, assessed as an increase in the magnitude of force fluctuations (i.e. coefficient of variation; Clark *et al.*, 2007). Importantly, maximal strength (Pijnappels *et al.*, 2008; Allet *et al.*, 2012) and the ability to control force (Davis *et al.*, 2021; Mear *et al.*, 2022) are significant predictors of balance and other ADLs (Enoka and Farina, 2021). It is no surprise, therefore, that both observational (Davis *et al.*, 2014; Willoughby and Copeland, 2015) and experimental (Caplan *et al.*, 2015) studies have demonstrated decreases in the ability to maintain balance with increased sedentary behaviour.

Fortunately, it has been suggested that brief, frequent muscular contractions throughout the day can have a potent influence on key physiological processes that mediate the adverse effects of prolonged sedentariness (Hamilton *et al.*, 2007; Dunstan *et al.*, 2012). Indeed, accumulating evidence suggests that interrupting long periods of sedentary time with regular physical activity “breaks” throughout the day has numerous health benefits, including increasing energy expenditure (Carter *et al.*, 2015), protecting cardiovascular endothelial function (Thosar *et al.*, 2015), lowering blood pressure (Larsen *et al.*, 2014), improving postprandial glucose metabolism (Bergouignan *et al.*, 2016) and decreasing lower back musculoskeletal discomfort (Thorp *et al.*, 2014). Furthermore, breaking up sedentary time is associated with decreased all-cause mortality (Katzmarzyk, 2014). These results suggest that relatively small changes in activity level and pattern have the potential to modify the adverse health risk of sedentary behaviour (Parry *et al.*, 2013). As such, minimising the amount of time spent in prolonged sitting and breaking up long periods of sitting as often as possible have now been included in governmental physical activity guidelines (Bull *et al.*, 2010; Brown *et al.*, 2012; World Health Organization, 2020). Nonetheless, the effects of breaking up sedentary time with physical activity breaks remains an understudied area, particularly in the context of neuromuscular function. Indeed, no study to date has investigated the effect of physical activity breaks on muscle strength, force control or balance.

Many studies on breaking up sedentary time have used either free walking or treadmill walking as their intervention (Latouche *et al.*, 2013; Larsen *et al.*, 2014; Thosar *et al.*, 2015; Bergouignan *et al.*, 2016). However, these are not always viable options, due to space and/or cost constraints (Carr *et al.*, 2012). Calisthenic exercises, such as body weight squats and lunges, represent an ideal alternative, in that they can be performed anywhere and without any equipment (Carter *et al.*, 2015). Previous research has demonstrated that breaking up short periods of sedentary behaviour (up to ~90 minutes) with calisthenics has a positive effect on energy expenditure and endothelial function (Carter *et al.*, 2015; Carter and Gladwell, 2017). A further advantage of calisthenics is that they target fitness components such as muscle strength and balance, which are key components of physical activity guidelines (Bull *et al.*, 2010) and are specifically affected by sedentary behaviour (Volkers *et al.*, 2012; Davis *et al.*, 2014). To our knowledge, the use of calisthenics as short-term intervention to break up sedentary time during the working day has not been studied.

The aim of the present study was to investigate the effect of breaking up sedentary time with regular bouts of calisthenic exercise on neuromuscular function. To that end, we compared the effects of a four-week calisthenics intervention with a control group who carried on with their normal behaviour. Neuromuscular function was assessed as muscle strength (maximal voluntary contraction of the knee extensors), muscle force control (measures of the magnitude and complexity of knee extensor force fluctuations) and dynamic balance (the Y balance test).

## Methods

### Participants

Healthy adults aged 18-65 who spent ≥6 hours a day sitting were recruited for the study. Sitting time was initially assessed via self-report, though was subsequently confirmed using accelerometery (activPAL4 micro; PAL Technologies Ltd., Glasgow, UK). 17 participants provided written informed consent to participate in the study, which was approved by the ethics committee of the University of Essex (Ref. ETH2021-0934), and which adhered to the Declaration of Helsinki. Participants were randomly assigned into a control group (n = 9) or a calisthenics intervention group (n = 8; see Table 1 for participant physical characteristics).

**Table 1.** Participant physical and sedentary behaviour characteristics. Values are means ± SD.

|  | Control (n = 9) | Intervention (n = 8) |
| --- | --- | --- |
| Age (years) | 29.9 ± 9.7 | 37.6 ± 15.8 |
| Height (m) | 1.74 ± 0.07 | 1.71 ± 0.12 |
| Body mass (kg) | 81.7 ± 22.7 | 71.6 ± 22.6 |
| Daily sitting time (hours) |  |  |
| Week 0 | 10.7 ± 1.7 | 9.1 ± 2.7 |
| Week 4 | 9.3 ± 1.4 | 10.3 ± 1.1 |
| Daily steps |  |  |
| Week 0 | 8354 ± 4919 | 7948 ± 3591 |
| Week 4 | 7759 ± 4068 | 8456 ± 3720 |

### Experimental design

Participants visited the laboratory on two separate occasions, five weeks apart. They were instructed to arrive at the laboratory in a rested state (having performed no strenuous exercise in the preceding 24 hours) and to have consumed neither food nor caffeinated beverages in the 3 hours prior to arrival. The first visit comprised familiarisation with testing equipment/experimental procedures and baseline (week 0) testing. Following this, all participants were instructed to carry on with their normal behaviour for the next week, so that baseline measures of sitting time could be confirmed using accelerometery. After this initial week, the control group were instructed to carry on with their normal behaviour for the next four weeks (weeks 1-4), while the calisthenics intervention group were given instructions on using calisthenics to break up sedentary time during the working week (i.e. 09:00-17:00, Monday-Friday) for the next four weeks (weeks 1-4; see *“Calisthenics intervention”* below for further details).

### Accelerometery

During their first visit to the laboratory, participants were provided with an ActivPAL4 physical activity logger (PAL Technologies Ltd., Glasgow, UK) to wear for the duration of the study. The ActivPAL4 is able to distinguish the incline of the leg to determine periods of lying, sitting, standing, and stepping (Harrington *et al.*, 2011). The ActivPAL4 unit was placed into a nitrile sleeve and then attached to participants left leg at the midline of the anterior thigh using a Tegaderm film dressing (3M Health Care, St Paul, Minnesota, USA). The nitrile sleeve and Tegaderm dressing providing a waterproof barrier, allowing participants to continue wearing the unit when bathing.

### Measures of neuromuscular function

On arrival at the laboratory, participants were assessed on the Y balance test, a measure of single leg dynamic balance (Plisky *et al.*, 2006). Assessment of dynamic balance was chosen given the dynamic nature of many ADLs (Ringhof and Stein, 2018). The Y balance test apparatus consists of an elevated central footplate (2.54 cm off the ground) and pipes, with reach indicator blocks, attached in the anterior, posteromedial, and posterolateral directions. Participants stood with one leg on the footplate, with the most distal aspect of their foot on a marked stating line. While maintaining single leg stance, participants then reached with their free leg in the anterior, posteromedial, and posterolateral directions (Plisky *et al.*, 2009). As a significant learning effect has previously been demonstrated (Hertel *et al.*, 2000), participants performed six practice trials in each of the three reach directions with each leg as the stance leg.

Following this practice, participants rested for 10 minutes, before performing three further attempts on each leg in which reach distance was recorded. A standardised testing order was used: participants started with their left leg as the stance leg and reached in the anterior, then posterolateral, and finally posteromedial directions. After doing this three times, participants repeated this with the right leg as the stance leg. All testing was conducted barefoot, to eliminate any additional balance and stability from the shoes (Coughlan *et al.*, 2012).

Following completion of the Y balance test, participants rested for 10 minutes. They were then seated in the chair of a Biodex System 4 isokinetic dynamometer (Biodex Medical Systems Inc., Shirley, New York, USA), initialised and calibrated according to the manufacturer’s instructions. Their right leg was attached to the lever arm of the dynamometer, with the seating position adjusted to ensure that the lateral epicondyle of the femur was in line with the axis of rotation of the lever arm. Participants sat with relative hip and knee angles of 85° and 90°, respectively, with full extension being 0°. The lower leg was securely attached to the lever arm above the malleoli with a padded Velcro strap, whilst further straps secured firmly across the waist and both shoulders prevented any extraneous movement and the use of the hip extensors during the isometric contractions. The isokinetic dynamometer was connected via a custom-built cable to a CED Micro 1401-4 (Cambridge Electronic Design, Cambridge, UK). Data were sampled at 1 kHz and collected in Spike2 (Version 10; Cambridge Electronic Design, Cambridge, UK).

To assess knee extensor muscle strength, participants performed a series of brief (3-second) isometric maximal voluntary contractions (MVCs), each of which were separated by 60-seconds rest. Participants were given a countdown, followed by strong verbal encouragement to maximise their effort. 10 minutes after the establishment of muscle strength, participants performed a series of targeted isometric knee extension contractions at 10, 20 and 40% MVC, respectively, to assess their ability to control submaximal force. These contraction intensities were chosen because they are typical of the demands of ADLs (Kern *et al.*, 2001). The targets were determined from the highest instantaneous force obtained during the baseline (week 0) MVCs. Participants performed three contractions at each intensity, with contractions held for 6-seconds and separated by 4-seconds rest (Pethick *et al.*, 2021; Mear *et al.*, 2022). The intensities were performed in a randomised order, with 2 minutes rest between each intensity. Participants were instructed to match their instantaneous force with a 1mm thick target bar superimposed on a display ~1m in front of them and were required to continue matching this target for as much of the 6-second contraction as possible.

### Calisthenics intervention

Participants in the calisthenics intervention group were provided with a calisthenics exercise programme to complete during weeks 1-4. The calisthenics programme consisted of five different exercises: squats, arm circles, calf raises, knees to opposite elbows, and lunges (Carter *et al.*, 2015; Carter and Gladwell, 2017). Prior to conducting the intervention, participants were provided with written and verbal instructions, and demonstrations of each exercise were given. They were then provided with the opportunity to practice any unfamiliar exercises.

Participants were instructed to perform the exercises during the working day, i.e. 09:00-17:00, Monday-Friday, using them to break up prolonged periods of sedentary time. In week 1, participants were instructed to perform 4 sets of the exercises per day. One set consisted of 8 repetitions of each exercise, at a rate of one repetition every three-seconds. Thus, each set of exercises took 2 minutes to complete. In week 2, participants were instructed to increase to 6 sets of the exercises per day and in weeks 3 and 4, participants were instructed to increase to 8 sets of the exercises per day. The reason for the progression in the number of sets across the intervention was twofold: 1) in order to conform to the progression principle of exercise training (Hoffman, 2002); and 2) pilot testing indicated that participants experienced delayed onset muscle soreness when performing 8 sets in the first week of the intervention, due to unaccustomed exercise, and this led to decreased adherence.

### Data analysis

The accelerometery data was analysed using the PAL software suite (Version 8; PAL Technologies Ltd., Glasgow, UK). Participants daily time spent sitting and daily step count were extracted and averaged across a five-day period (Monday-Friday). Technical issues meant that baseline data were only available for 7 participants in the control group and 6 participants in the intervention group.

For the Y balance test, the greatest of the three trials was used for analysis of reach distance in each direction. Reach distance is significantly correlated with leg length (Gribble and Hertel, 2003). As such, reach distance was normalised to leg length, measured as the distance in centimetres from the anterior superior iliac spine to the centre of the ipsilateral medial malleolus. The normalised value was calculated as: (reach distance/leg length) x 100. The normalised reach distance was, therefore, expressed as a percentage of leg length. The greatest normalised reach distance from each direction was also summed to yield a composite reach distance. Composite reach was calculated as: (sum of three reach directions/three times leg length) x 100.

Maximal muscle strength was determined as the highest instantaneous force obtained during the MVCs. For the submaximal force control tasks, the mean value of the three contractions at each intensity was calculated. Values for individual contractions were calculated based on the steadiest 5 seconds of each contraction, with MATLAB (R2017a; The MathWorks, Massachusetts, USA) code identifying the 5 seconds of each contraction with the lowest standard deviation (SD). It has been recommended that both magnitude- and complexity-based measures should be used when characterising force control (Pethick *et al.*, 2021a). Magnitude-based measures of force control characterise force steadiness, while complexity-based measures reflect force adaptability; that is, the ability to modulate force output rapidly and accurately in response to task demands (Pethick *et al.*, 2021b).

The magnitude of variability in force output was assessed using the SD and coefficient of variation (CV), which provide measures of the absolute amount of variability in an output and the amount of variability normalised to the mean of the output, respectively (Enoka *et al.*, 2003). The complexity of force output was assessed using approximate entropy (ApEn) and detrended fluctuation analysis (DFA) α. ApEn was used to assess the regularity or randomness of force output (Pincus, 1991), while DFA was used to estimate temporal fractal scaling. The calculations of ApEn and DFA are detailed in Pethick *et al.* (2015). In brief, ApEn was calculated with template length, *m,* set at 2 and the tolerance for accepting matches, *r,* set at 10% of the SD of force output, and DFA was calculated across time scales (57 boxes ranging from 1250 to 4 data points).

### Statistics

All data are presented as means ± SD. Results were deemed statistically significant when *P* < 0.05. All data were tested for normality using the Shapiro-Wilk test. Independent samples *t*-tests were used to compare the physical characteristics (age, height, body mass) and baseline sedentary behaviour (sitting time, step count) of the control and calisthenics intervention groups. Student’s paired samples *t*-tests were used to compare the values for muscle strength, muscle force control (SD, CV, ApEn and DFA at 10, 20 and 40% MVC) and dynamic balance (anterior, posteromedial, posterolateral and composite reach in each leg) obtained at baseline and in week 4 in the control and calisthenics intervention groups.

## Results

### Participant physical and sedentary behaviour characteristics

The physical and sedentary behaviour characteristics of the control and intervention groups are presented in Table 1. There were no significant differences between the control and calisthenics intervention groups for age (*P* = 0.237), height (*P* = 0.521), body mass (*P* = 0.370), baseline daily sitting time (*P* = 0.209) or baseline daily step count (*P* = 0.870). All participants for whom ActivPAL data were available met the criteria of at least 6 hours a day sitting time.

### Sedentary behaviour

There were no changes from week 0 to week 4 in either the calisthenics intervention group or control group for daily sitting time (*P* = 0.332, *P* = 0.146, respectively) or step count (*P* = 0.814, *P* = 0.980, respectively; Table 1).

### Muscle strength

There was a significant increase in knee extensor MVC from week 0 to week 4 in the calisthenics intervention group (*P* = 0.036) but not the control group (*P* = 0.710; Figure 1; Table 2).

**Figure 1.**
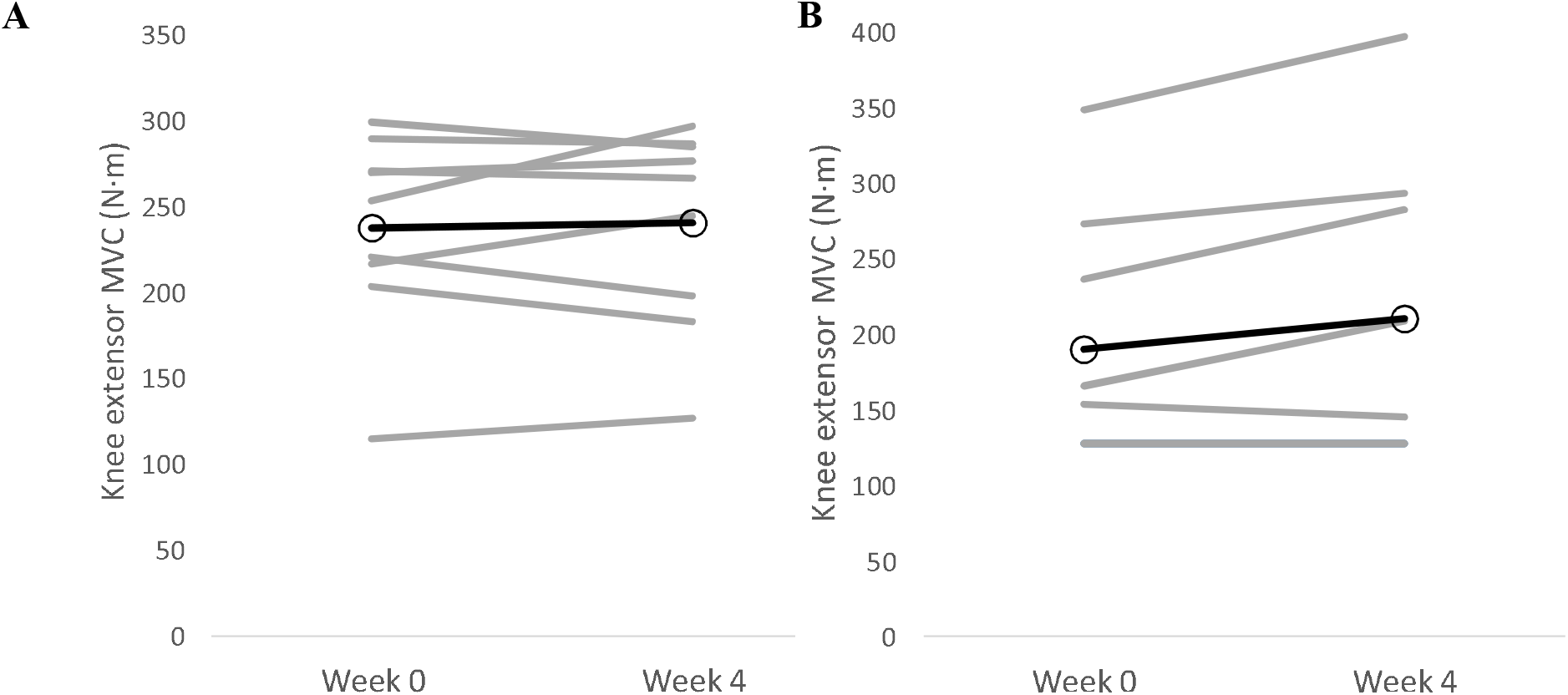
Individual changes (grey lines) in knee extensor MVC strength from baseline (week 0) to the end of the intervention (week 4). Panel A, control group; Panel B, calisthenics intervention group. Dark lines represent group means.

**Table 2.**
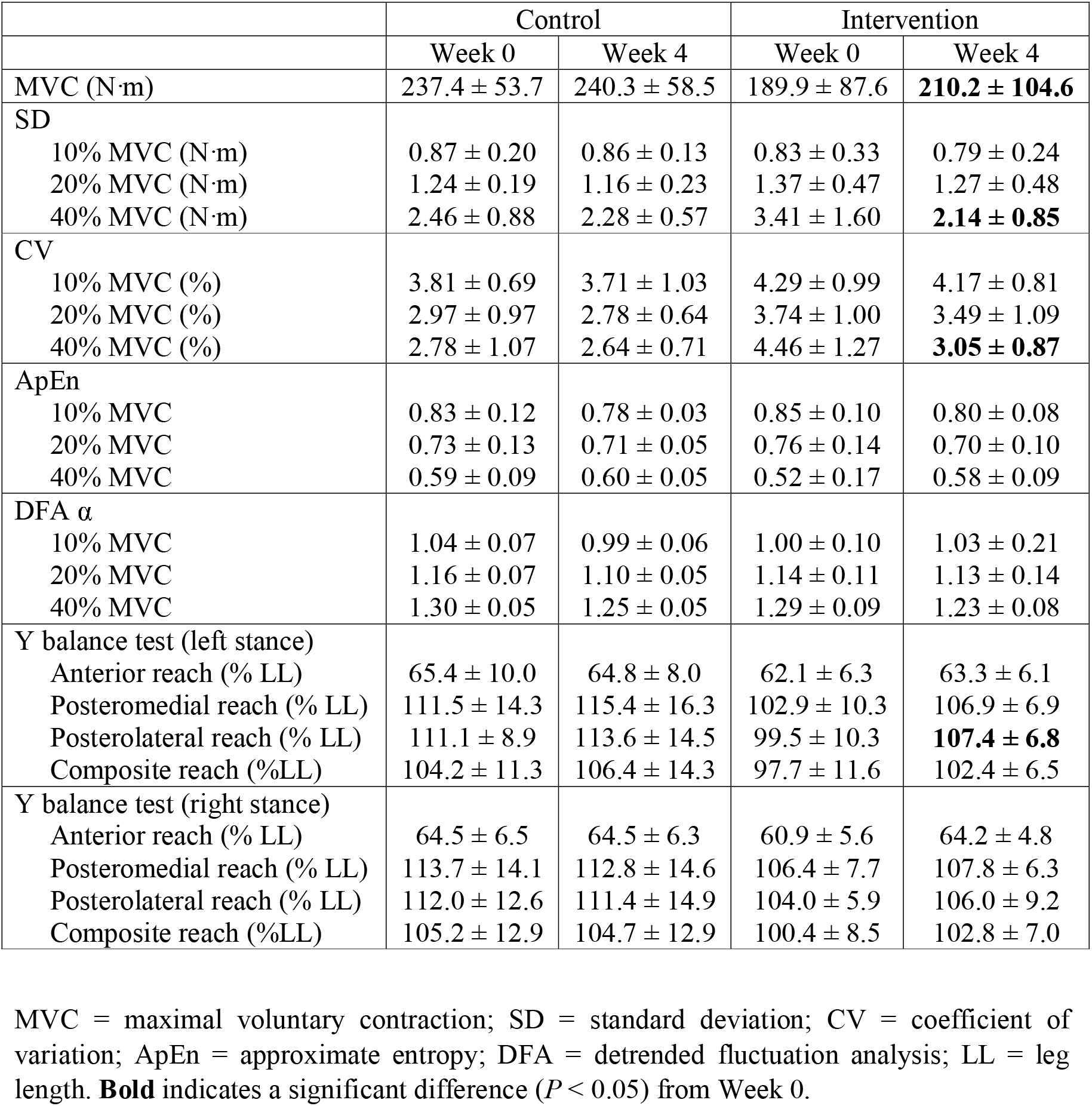
Knee extensor strength, force control and Y balance test data from baseline (week 0) and the end of the intervention (week 4).

|  | Control |  | Intervention |  |
| --- | --- | --- | --- | --- |
|  | Week 0 | Week 4 | Week 0 | Week 4 |
| MVC (N·m) | 237.4 ± 53.7 | 240.3 ± 58.5 | 189.9 ± 87.6 | **210.2 ± 104.6** |
| SD |  |  |  |  |
| 10% MVC (N·m) | 0.87 ± 0.20 | 0.86 ± 0.13 | 0.83 ± 0.33 | 0.79 ± 0.24 |
| 20% MVC (N·m) | 1.24 ± 0.19 | 1.16 ± 0.23 | 1.37 ± 0.47 | 1.27 ± 0.48 |
| 40% MVC (N·m) | 2.46 ± 0.88 | 2.28 ± 0.57 | 3.41 ± 1.60 | **2.14 ± 0.85** |
| CV |  |  |  |  |
| 10% MVC (%) | 3.81 ± 0.69 | 3.71 ± 1.03 | 4.29 ± 0.99 | 4.17 ± 0.81 |
| 20% MVC (%) | 2.97 ± 0.97 | 2.78 ± 0.64 | 3.74 ± 1.00 | 3.49 ± 1.09 |
| 40% MVC (%) | 2.78 ± 1.07 | 2.64 ± 0.71 | 4.46 ± 1.27 | **3.05 ± 0.87** |
| ApEn |  |  |  |  |
| 10% MVC | 0.83 ± 0.12 | 0.78 ± 0.03 | 0.85 ± 0.10 | 0.80 ± 0.08 |
| 20% MVC | 0.73 ± 0.13 | 0.71 ± 0.05 | 0.76 ± 0.14 | 0.70 ± 0.10 |
| 40% MVC | 0.59 ± 0.09 | 0.60 ± 0.05 | 0.52 ± 0.17 | 0.58 ± 0.09 |
| DFA $\alpha$ | | | | |
| 10% MVC | 1.04 ± 0.07 | 0.99 ± 0.06 | 1.00 ± 0.10 | 1.03 ± 0.21 |
| 20% MVC | 1.16 ± 0.07 | 1.10 ± 0.05 | 1.14 ± 0.11 | 1.13 ± 0.14 |
| 40% MVC | 1.30 ± 0.05 | 1.25 ± 0.05 | 1.29 ± 0.09 | 1.23 ± 0.08 |
| Y balance test (left stance) |  |  |  |  |
| Anterior reach (% LL) | 65.4 ± 10.0 | 64.8 ± 8.0 | 62.1 ± 6.3 | 63.3 ± 6.1 |
| Posteromedial reach (% LL) | 111.5 ± 14.3 | 115.4 ± 16.3 | 102.9 ± 10.3 | 106.9 ± 6.9 |
| Posterolateral reach (% LL) | 111.1 ± 8.9 | 113.6 ± 14.5 | 99.5 ± 10.3 | **107.4 ± 6.8** |
| Composite reach (%LL) | 104.2 ± 11.3 | 106.4 ± 14.3 | 97.7 ± 11.6 | 102.4 ± 6.5 |
| Y balance test (right stance) |  |  |  |  |
| Anterior reach (% LL) | 64.5 ± 6.5 | 64.5 ± 6.3 | 60.9 ± 5.6 | 64.2 ± 4.8 |
| Posteromedial reach (% LL) | 113.7 ± 14.1 | 112.8 ± 14.6 | 106.4 ± 7.7 | 107.8 ± 6.3 |
| Posterolateral reach (% LL) | 112.0 ± 12.6 | 111.4 ± 14.9 | 104.0 ± 5.9 | 106.0 ± 9.2 |
| Composite reach (%LL) | 105.2 ± 12.9 | 104.7 ± 12.9 | 100.4 ± 8.5 | 102.8 ± 7.0 |

MVC = maximal voluntary contraction; SD = standard deviation; CV = coefficient of variation; ApEn = approximate entropy; DFA = detrended fluctuation analysis; LL = leg length. **Bold** indicates a significant difference ( $P < 0.05$ ) from Week 0.

### Muscle force control

The knee extensor force control data from weeks 0 and 4 are presented in Table 2. There were no changes in any of the force control measures during contractions at 10% MVC from week 0 to week 4 in the calisthenics intervention group (SD: *P* = 0.571; CV: *P* = 0.658; ApEn: *P* = 0.750; DFA α: *P* = 0.638). Similarly, there were no changes in any of the force control measures during contractions at 20% MVC from week 0 to week 4 in the calisthenics intervention group (SD: *P* = 0.208; CV: *P* = 0.147; ApEn: *P* = 0.077; DFA α: *P* = 0.625). There were significant decreases in SD (*P* = 0.031) and CV (*P* = 0.016) during contractions at 40% MVC from week 0 to week 4 in the calisthenics intervention group (Figure 2; Table 2). There were no changes in ApEn (*P* = 0.283) or DFA α (*P* = 0.176) during contractions at 40% MVC in the calisthenics intervention group.

**Figure 2.**
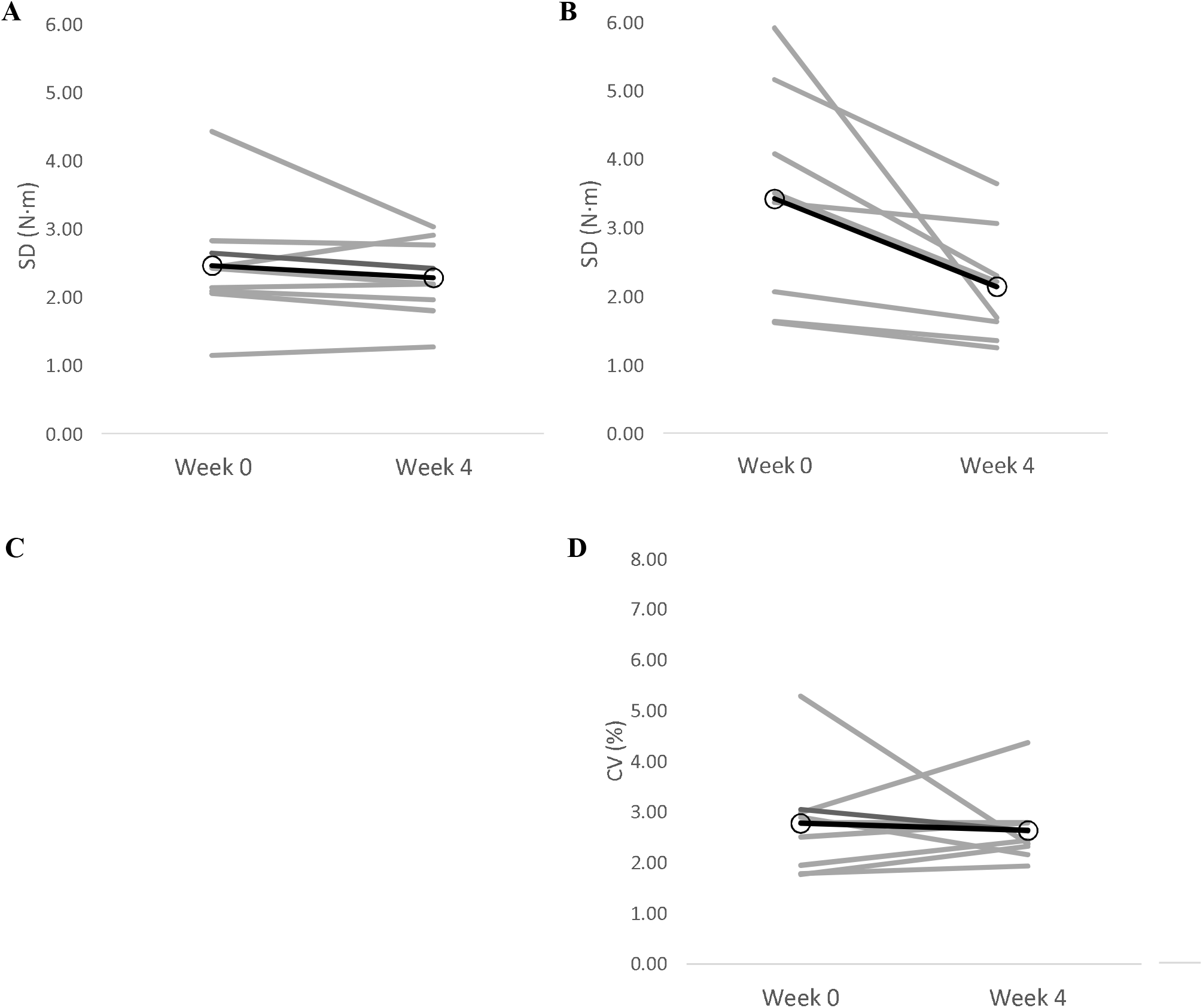
Individual changes (grey lines) in knee extensor force control from baseline (week 0) to the end of the intervention (week 4). Panel A, control group SD; Panel B, calisthenics intervention group SD; Panel C, control group CV; Panel D, calisthenics intervention group CV. Dark lines represent group means.

There were no changes in any of the force control measures during contractions at 10% MVC (SD: *P* = 0.816; CV: *P* = 0.786; ApEn: *P* = 0.206; DFA α: *P* = 0.133), 20% MVC (SD: *P* = 0.442; CV: *P* = 0.451; ApEn: *P* = 0.631; DFA α: *P* = 0.156) or 40% MVC (SD: *P* = 0.303; CV: *P* = 0.740; ApEn: *P* = 0.286; DFA α: *P* = 0.094) from week 0 to week 4 in the control group.

### Dynamic balance

The Y balance test data from weeks 0 and 4 are presented in Table 2. In the calisthenics intervention, group there was a significant increase in Y balance test normalised posterolateral reach distance with left leg stance from week 0 to week 4 (*P* = 0.046; Figure 3). There were no changes in normalised anterior (*P* = 0.447), posteromedial (*P* = 0.210) or composite (*P* = 0.069) reach distance in the calisthenics intervention group. There were no changes in normalised reach distance in any direction with the right leg from week 0 to week 4 in the calisthenics intervention group (anterior: *P* = 0.092; posteromedial: *P* = 0.631; posterolateral: *P* = 0.479; composite: *P* = 0.159).

**Figure 3.**
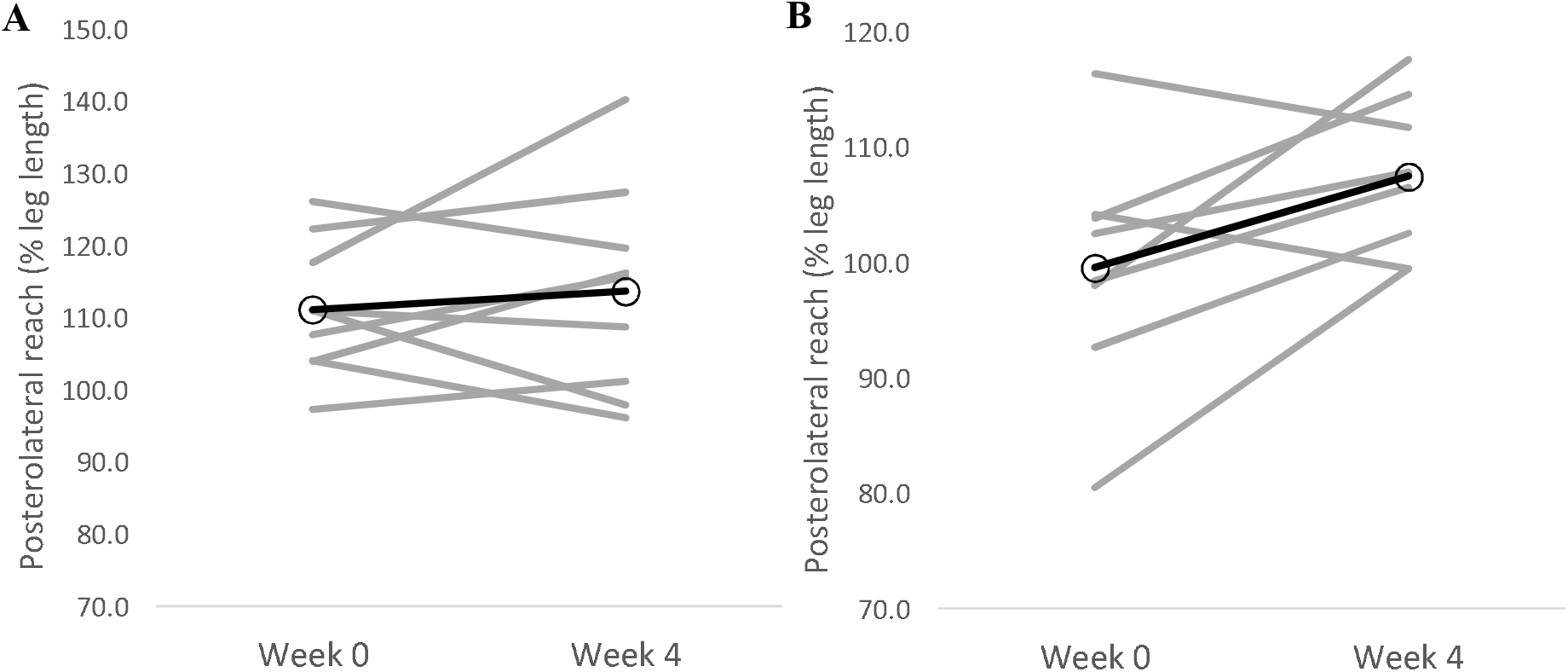
Individual changes (grey lines) in Y balance test posterolateral reach with left leg stance from baseline (week 0) to the end of the intervention (week 4). Panel A, control group; Panel B, calisthenics intervention group CV. Dark lines represent group means.

In the control group, there were no changes in normalised reach distance in any direction with either the left (anterior: *P* = 0.687; posteromedial: *P* = 0.329; posterolateral: *P* = 0.532; composite: *P* = 0.450) or right (anterior: *P* = 0.988; posteromedial: *P* = 0.799; posterolateral: *P* = 0.682; composite: *P* = 0.780) legs from week 0 to week 4.

## Discussion

The major novel finding of the present study was that a short term (4-week) intervention designed to break up sedentary time with calisthenics exercise was effective at improving neuromuscular function. Specifically, the calisthenics intervention increased knee extensor MVC, knee extensor force steadiness (SD and CV) during contractions at 40% MVC and posterolateral reach with left leg stance in the Y balance test, whereas no such changes were evident in the control group. These results indicate the potential of calisthenics, i.e. simple body weight exercises that can be performed anywhere and without any equipment, as a tool to break up sedentary time and mitigate against its deleterious functional consequences.

### Effect of calisthenics on neuromuscular function

The present study adds to the growing body of literature demonstrating the positive physiological effects of interventions designed to break up sedentary time with physical activity (Benatti and Ried-Larsen, 2015; del Pozo-Cruz *et al.*, 2018). It is the first experimental study to demonstrate that regularly breaking up sedentary time with calisthenics has a positive effect on objective measures of neuromuscular function (i.e. muscle strength, muscle force control and balance; Figures 1-3; Table 2). This is in line with previous cross-sectional studies demonstrating better performance of ADLs with more frequent breaks in sedentary time (Sardinha *et al.*, 2015). Moreover, the presently observed improvements were evident with only 16 minutes per day of calisthenics exercise, spread across a period of 8 hours; thus providing evidence that small changes in activity level and pattern are able to modify neuromuscular function. Indeed, the daily requirement of the calisthenics intervention was such that it altered neither participants daily sitting time nor step count (Table 2). These results illustrate the utility of calisthenics as an intervention to break up sedentary time and improve neuromuscular function and serve to highlight their potential to improve other aspects of physiological function. Calisthenics exercises can activate both upper and lower limb musculature, and target multiple components of fitness (Carter *et al.*, 2015; Carter and Gladwell, 2017), making them more effective than previously used interventions such as free walking, treadmill walking or portable pedal machines; can be performed anywhere, including in the workplace or at home (Carter *et al.*, 2015); and do not require any costly or space consuming equipment (Carr *et al.*, 2012).

The results of the present study demonstrated that using calisthenics to break up sedentary time throughout the day can improve knee extensor strength (Figure 1; Table 2). The effectiveness of the calisthenics intervention was likely due to the squat and lunge exercises, both of which target the knee extensors (Jönhagen *et al.*, 2009; Slater and Hart, 2017). Though no such measures were taken in the present study, previous research found no change in muscle thickness, indicative of hypertrophy, following 4 weeks of upper body calisthenics training (Kotarsky *et al.*, 2018). As such, it is reasonable to suggest that the presently observed increase in strength was not due to muscle hypertrophy. Rather, it was likely mediated by decreases in motor unit recruitment threshold and increases in motor unit discharge rate (Del Vecchio *et al.*, 2019). Notably, decreased motor unit firing rates have been observed with enforced periods of inactivity (Duchateau and Hainaut, 1990; Seki *et al.*, 2001). The observed increase in strength has important health and functional implications. Lower extremity strength has been demonstrated to be predictive of all-cause mortality (Li *et al.*, 2018). Indeed, for every 15 N increase in strength (similar to that observed in the present study; Table 2), there is a 7% reduced risk of all-cause mortality (Loprinzi, 2016). Knee extensor strength is also an important predictor of performance in ADLs, including balance (Allet *et al.*, 2012) and locomotion (Kang and Dingwell, 2008).

The calisthenics intervention resulted in a significant decrease in the magnitude of knee extensor force fluctuations, as measured by the SD and CV, during contractions at 40% MVC (Figure 2; Table 2). These changes are indicative of increased force steadiness (Pethick *et al.*, 2021b). In contrast, the intervention had no effect on the complexity of force output, as measured by ApEn and DFA α, indicating no change in force adaptability (Pethick *et al.*, 2021b). The increase in force steadiness at 40% but not 10 or 20% MVC could be due to the use of gross motor movements (i.e. squats and lunges) that produce large overall forces (Keogh *et al.*, 2007). Knee extensor EMG activity during the squat, for example, has been demonstrated to reach up to 63% of the value obtained during an isometric MVC (Isear Jr. *et al.*, 1997). Muscle force CV is strongly associated with variance in common synaptic input to motor neurons (Enoka and Farina, 2021). Thus, as with the increase in strength, it is likely that the increase in force steadiness was mediated an increase in motor neuron output, specifically an increase in motor unit discharge rate (Del Vecchio *et al.*, 2019). The intervention-induced increase in knee extensor force steadiness has the potential to be as, if not more, important than the increase in strength. Indeed, previous research has demonstrated that force control is a stronger predictor of both static (Davis *et al*., 2020) and dynamic balance (Mear *et al*., 2022) than maximal strength.

Interestingly, Y balance test normalised posterolateral reach with left leg stance, but not anterior or posteromedial reach, increased as a result of the calisthenics intervention (Figure 3; Table 2). This increase in posterolateral reach is indicative of greater dynamic balance and stability (Lockie *et al.*, 2013). The calisthenics exercises were, for the most part, linear exercises performed in the sagittal plane. Consequently, anterior reach, also performed in the sagittal plane, would have been expected to be the most likely of the reach directions to improve. Why only posterolateral reach increased as a result of the intervention is unclear, though it must be noted that previous training studies have often failed to demonstrate increases in all reach directions (Benis *et al.*, 2016; Xiong *et al.*, 2022). No improvements in reach with right leg stance were observed as a result of the intervention (Table 2). With the exception of anterior reach, the right leg was the better performing leg at week 0 (Table 2). Thus, the improvement in the left leg, but not the right leg, could simply be reflective of the left leg having greater scope to improve.

### Limitations and suggestions for future research

The present study was subject to several limitations, which can be broadly categorised as relating to the intervention, the population tested and the study design. With regards to the intervention, it consisted of squats, arm circles, calf raises, knees to opposite elbows, and lunges, as previously used by Carter *et al.* (2015). Given the specificity principle of training, it could be argued that arm circles are unlikely to contribute to increases in knee extensor maximal strength, force control or dynamic balance. As such, future studies using calisthenics to break up sedentary and improve neuromuscular function should ensure that all exercises used are specific to the outcomes being measured. Future studies could also manipulate the intensity (number of sets and repetitions) and/or duration of the intervention, thus conforming to the progression principle of training. Higher intensity or longer duration studies may be necessary to see improvements in the variables that did not change in the present study.

With regards to the population tested, our participants comprised a mix of university staff and students, who (ActivPAL data indicated) exhibited markedly different daily patterns of activity. It may be of utility to investigate more homogeneous populations, such as office workers, particularly given that sedentary behaviour can account for >80% of work time (Parry and Straker, 2013). Indeed, given its simplicity to conduct, calisthenics has been suggested to be an ideal workplace intervention (Carter and Gladwell, 2017).

Because the purpose of the study was simply to ascertain the effectiveness of breaking up sedentary time with calisthenics exercise on neuromuscular function, we did not take any measures that could determine the mechanism(s) for the intervention-induced improvements. Future studies could use techniques such as high-density electromyography to assess neural effects (e.g. motor unit recruitment and discharge properties) and imaging modalities such dual-energy x-ray absorptiometry to assess muscular effects (e.g. change in body composition/body size). Furthermore, we did not follow up with participants again beyond the end of the intervention. As such, it is unclear for how long the intervention-induced improvements in strength, force control and dynamic balance were retained.

### Conclusion

Sedentary behaviour is associated with decreased maximal strength, decreased ability to control force and decreased ability to maintain balance. The present findings are the first to demonstrate that breaking up sedentary time with calisthenics exercise has a positive effect on these important indices of neuromuscular function. Calisthenics exercise overcomes the limitations of other physical activity interventions to break up sedentary time, in that it can be performed anywhere, without any equipment and targets multiple fitness components. Importantly, these results indicate that small changes in activity level and pattern (only 16 minutes per day, spread across 8 hours) can mitigate against the negative effects of prolonged sedentary time.

## Author contribution

The experiments were performed at the University of Essex. JP and VG conceptualised the study. JP and EM acquired the data. All authors were involved in analysing and interpreting the data. All authors contributed to drafting and critically revising the manuscript.

